# The evolutionary potential of diet-dependent effects on lifespan and fecundity in a multi-parental population of *Drosophila melanogaster*

**DOI:** 10.1101/343947

**Authors:** Enoch Ng’oma, Wilton Fidelis, Kevin M. Middleton, Elizabeth G. King

## Abstract

The nutritional conditions experienced by a population play a major role in shaping trait evolution in many taxa. Constraints exerted by nutrient limitation or nutrient imbalance can influence the maximal value that fitness components such as reproduction and lifespan attains, and organisms may shift how resources are allocated to different structures and functions in response to changes in nutrition. Whether the phenotypic changes associated with changes in nutrition represent an adaptive response is largely unknown. Further, it is unclear whether the response of fitness components to diet even has the potential to evolve in most systems. In this study, we use an admixed multiparental population of *Drosophila melanogaster* reared in three different diet conditions to estimate quantitative genetic parameters for lifespan and fecundity. We find significant genetic variation for both traits in our population and show that lifespan has moderate to high heritabilities within diets. Genetic correlations for lifespan between diets were significantly less than one, demonstrating a strong genotype by diet interaction. These findings demonstrate substantial standing genetic variation in our population that is comparable to natural populations and highlights the potential for adaptation to changing nutritional environments.

## Introduction

An organism’s diet has a direct influence on nearly all phenotypes by determining the amount of nutrients and energy available to build structures and perform functions. In addition to this unavoidable direct effect, organisms may also alter how resources are allocated to different traits when experiencing different diets. In an environment where resource availability varies across space and/or time, and the optimal allocation of resources changes with diet, we expect to see the evolution of such phenotypically plastic resource allocation strategies.

In many non-human populations including nematode (Greer and Brunet, 2009), fruit fly (Chippindale *et al.*, 1993b), killifish (Vrtílek and Reichard, 2014), waterstrider (Kaitala, 1991) and mouse (Sprott, 1997), nutrient limitation without starvation, hereafter referred to as dietary restriction (DR), results in lifespan extension, and is often but not always accompanied by a reduction in reproductive output (Hsin and Kenyon, 1999; Barnes *et al.*, 2006; Flatt *et al.*, 2008). These diet-dependent changes have been hypothesized to result from an increased allocation of a limited nutritional resource to somatic maintenance (van Noordwijk and de Jong, 1986; Sohal and Weindruch, 1996; Simmons and Bradley, 1997; Hughes and Reynolds, 2005; Kirkwood, 1977; Shanely and Kirkwood, 2000), and decreased allocation to reproduction. Previous models predict this pattern is adaptive in environments with fluctuating resource availability (Neel, 1962; Wells, 2009; Fisher *et al.*, 2010; Speakman 2011; Warren *et al.*, 2013). However, the main physiological and evolutionary drivers of the DR effect and its underlying mechanisms are still not well-understood. Several studies taking a nutritional geometry approach, which measure phenotypes across serial dilutions of diet components, have found that lifespan and reproduction are affected primarily by the protein-carbohydrate ratio of the diet rather than the caloric content (e.g. Skorupa *et al*, 2008; Lee *et al*, 2008; Jensen *et al*, 2015). These studies demonstrate that calorie limitation *per se*, is not driving these phenotypic patterns, though it may be that individual diet components act as a cue that induces an optimal allocation of resources in wild populations. In addition, recent meta-analyses quantifying the nutritional composition of the various types of diets that are used in DR studies have shown that DR has greater effect on reproduction and lifespan in model species than it has on non-model species, suggesting the effect may be due to laboratory adaptation (Speakman, 2011; Nakagawa *et al*, 2012, Moatt *et al*, 2016). However, different hypotheses for the evolution of the DR response have rarely been formally tested (but see Zajitschek *et al,* 2016; 2018), nor has the evolutionary potential of the focal traits been established in most systems (Ng’oma *et al.*, 2017), which is an important first step.

The capacity for a given trait to evolve in a population depends on the proportion of trait variation that is attributable to genetic variation and the intensity of selection acting on it (Roff 2006; Brčić-Kostić, 2010). Selection can have a direct effect on trait evolution if the trait directly affects fitness, or indirectly if the trait is correlated with a trait under selection (Lande and Arnold, 1983; Hunt *et al.*, 2004). However, even when a selective pressure is observed to act on the trait, the amount of evolutionary change depends on the amount of genetic variation expressed for that trait (Fisher, 1930). Quantitative genetic parameters, such as the heritability of a given trait, remain some of the more informative parameters we can estimate in order to learn about the genetic basis of a trait, to predict trait values in a population, and to predict the selection response, even with the advances in genomics and molecular genetics in recent years. Recent experimental evolution studies exposing fruit flies to DR have demonstrated that lifespan and reproduction do respond to selection under different diet conditions, although lifespan appears to be decoupled from fecundity (Zajitschek *et al,* 2016; 2018). In addition, studies in many species have frequently demonstrated genotype by diet variation for complex traits, such as gene expression and fitness in yeast (Gagneur *et al.*, 2013), cuticular hydrocarbons in *D. simulans* (Ingleby *et al.*, 2013), and metabolic phenotypes in *D. melanogaster* (Reed *et al.*, 2010, 2014; Qi *et al.*, 2012). These studies suggest that genotype by environment interactions (GEI) are common and could be important in the nutrition-dependent response of life history traits such as fecundity and lifespan.

There has been a significant effort to understand the genetic basis for the dietary restriction (DR) response with both molecular genetic approaches (Mair and Dillin, 2008; Grandison *et al.*, 2009; Teleman, 2010; Nässel *et al.*, 2015) and mapping approaches (Rikke *et al.*, 2010; Stanley *et al.*, 2017). As is the case for most complex traits, mapping approaches to identify the genetic loci determining how lifespan changes with diet have failed to identify the specific contributors. Studies focused on mapping in natural populations often do not identify the regions of the genome as QTL that harbor candidate genes identified by molecular approaches (Remolina *et al.*, 2012; Burke, King, *et al.*, 2014; Reed *et al.*, 2014; Carnes *et al.*, 2015) and there are several studies in which gene expression patterns do not change in the direction one would expect from these loss-of-function studies (Min *et al.*, 2008; Giannakou *et al.*, 2008; Stanley *et al.*, 2017). In light of these studies, a possible hypothesis is that the DR response is highly polygenic and best understood by quantifying holistic measures of genetic variation, such as the heritability and GEI. Additionally, it is possible that the limited genetic variation included in QTL mapping approaches limits the ability to identify causative variants that are segregating in natural populations. The majority of QTL studies in the past have used only two genomes to create the mapping population. More recently, multiparent populations (MPPs) have been gaining in popularity, where multiple (typically at least 4, and often many more) genomes are used to seed the population, thereby including increased genetic variability (Kover *et al.*, 2009; McMullen *et al.*, 2009; Huang *et al.*, 2011; Aylor *et al.*, 2011; Threadgill and Churchill, 2012; King, *et al.*, 2012; King, *et al.*, 2012; Cubillos *et al.*, 2013).

In this paper, we employ a half-sibling, split environment design using an admixed multiparent population derived from an established mapping resource, the *Drosophila* Synthetic Population Resource (DSPR), to estimate quantitative genetic parameters for lifespan and fecundity in three different diets. Our diets include typical DR and control diets used in *D. melanogaster* following Bass *et al* (2007), and a high sugar diet that contains ∼7 times the amount of sugar relative to our control diet. We are able to estimate the evolutionary potential of how lifespan and fecundity respond to these diets in order to establish, 1) the ability of these traits to evolve in general, and provide specific estimates of quantitative genetic parameters, 2) demonstrate levels of genetic variation that are comparable with wild derived populations, and 3) establish the validity of the evolve and resequence approach using a synthetic population as a base population. We expect to find extended lifespan and reduced fecundity on the DR diet relative to the control and reduced lifespan and fecundity in the high sugar diet relative to the control. More importantly, however, we expect to see substantial variablity among families in the phenotypic patterns in response to our diet treatments, with estimates of heritabilities within diets and genetic correlations between diets showing evidence for substantial genetic variation.

## Methods

### Experimental setup

*Generation of experimental population.* We used an eight-way multi-parent population, the DSPR, to characterize the phenotypic response of lifespan and fecundity of mated female *D. melanogaster* to three nutritional environments and to estimate the heritabilities of these traits. The development, and genetic properties of the DSPR are described in detail in King *et al.*, (2012a) and King *et al.*, (2012b). Briefly, the DSPR is a set of recombinant inbred lines created from 8 inbred founder lines that were crossed for 50 generations and then inbred via full sibling mating for 25 generations. To generate our starting population, for each of 835 recombinant inbred lines (RILs) of the *B* sub-population of the DSPR, five young females that had mated intra-line in vials, were randomly selected and distributed to each of six cages (20.3 cm × 20.3 cm × 20.3 cm). Eggs were collected in milk bottles (250 mL) twice on the next two successive days, each after a 22-hour oviposition period on maintenance diet (Table S1). Eclosed flies were released to six cages in populations >2,000 per cage and allowed to mate *en-masse* for five generations. A polystyrene plate, 100 mm × 15 mm (cat. No. FB0875713 Fisher Scientific, USA) containing maintenance food was supplied to each cage three times a week (Monday, Wednesday, Friday). A separate micro-plate (60 mm × 15 mm, cat. no. FB0875713A, Fisher Scientific, USA) with moist cotton wool was supplied to each cage to serve as a supplemental drinking source. Very thin slices of food with visually estimated 50-90 eggs were transferred to 30 vials (25 mm × 95 mm, Polystyrene Reload, cat. no. 32-109RL, Genesee Scientific, USA) for each cage for each successive generation. Egg vials from each cage for each next generation were distributed across all cages to ensure a genetically homogenous experimental population. We maintained all populations on a three week cycle (21 days from oviposition to egg collection) on a cornmeal-dextrose-yeast diet for 5 generations before the half-sibling experiment was started.

*Experimental design.* We employed a split-environment, half-sibling breeding design to determine the phenotypic effects of three dietary conditions and estimate quantitative genetic parameters. We chose to use a dietary restriction and control diet similar to those commonly used in studies of the lifespan effects of DR in flies (Bass et al. 2007). We also employed a high sugar diet, with the same amount of yeast as the control diet but with ∼7 times the amount of sugar. The motivation for choosing this diet was to provide a high calorie diet that has the potential to serve as a model of some modern human diets (i.e. a higher calorie, higher carbohydrate diet than the diet experienced for the majority of evolutionary history). While it would have been ideal to measure a complete range of diets with many possible combinations of yeast and sugar, the scale of this experiment limited the number of diets that could be examined. We created a set of paternal half-sibling families by sequentially mating each of 28 males to three females (here on, sires and dams, respectively) on a common cornmeal-dextrose-yeast (maintenance) diet (Table S1). Each dam was housed in its own vial and each sire rotated among these three vials. In round one, to ensure each dam had a mating opportunity early, the sire spent 2 days with each dam. Each sire was then rotated among the three dams for two additional rounds lasting 3.5 days each encounter. Two dams were kept waiting in media vials when the sire was matched with the third dam. Whole families were restarted with new flies when a sire died during crossing. In case of death of a waiting unmated dam, only that dam was replaced, whereas if a mated dam died, a new dam family was restarted. Starting with 61 sires and 183 dams, 28 half-sibling families and 77 maternal full-sibling families successfully produced enough offspring to be included in the experiment (Fig. 1).

**Figure 1:**
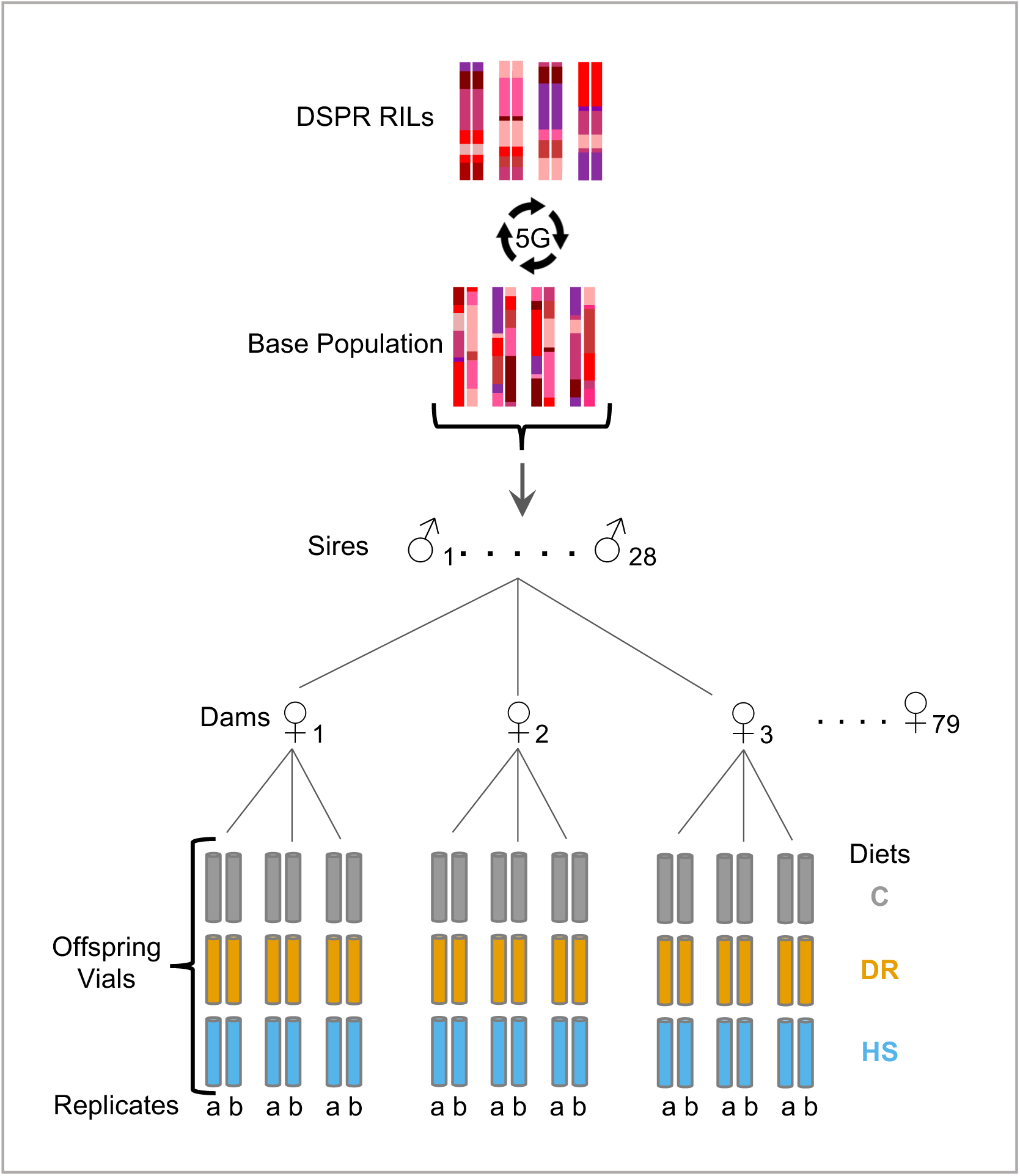
Experimental design. A set of 835 DSPR RILs were crossed for 5 generations to create an outbred synthetic population. From this population, 28 randomly chosen sires were each mated to 3 dams (actual N = 77 ♀). The resulting offspring from each dam were split to each of C, DR, and HS diet treatments with two replicate vials in each diet.

From each of the 77 dams, in most cases, we set up two replicate vials of 24 female and 6 male offspring in each of three diet treatments (described below). We included males to provide mating opportunities to females throughout their lifetime but chose a smaller number of males to reduce harassment of females within the vials. As expected, even successful sire and dam pairs did not always produce enough offspring for a full complement. Thus, some vials contained fewer than 24 females though never below 15 females, and the dataset is not fully balanced with all sire families having all three dams, split in all three diets, and with two replicates per diet. The realized totals of sire half-sibling families and dam full-sibling families available for analysis in each diet are presented in Table S2.

*Diet treatments.* Offspring of each sire were split across three experimental diets: control (C), dietary restriction (DR), and high sugar (HS). These diets were also used in our previous mapping study using the DSPR RILs by Stanley *et al*. (2017) and the composition of each is detailed in Table S1. We used the SAFPro Relax + YF 73050 brand of yeast typically containing 45-60g of protein and 30-38g of carbohydrate per 100g inactivated yeast (Lesaffre Yeast Corp., Milwaukee, USA). To preserve quality, diets were stored at 4 °C and used within two weeks of preparation. To permit phenotype measurement and ensure that food did not degrade, individuals were moved to vials with fresh food three times per week. All flies in all experiments described here were reared in a growth chamber at 23° C, ≥ 50% relative humidity, and a 24:0 light:dark cycle, which are the typical maintenance conditions for the DSPR flies.

### Phenotype measurement

We measured two phenotypes: 1) lifespan - the total number of days each fly lived from the date of eclosion, and 2) weekly fecundity - the total number of eggs a set of females in each vial produced within a 24-hour period measured once per week. Both phenotypes were measured from the same set of experimental vials over a full lifespan. For lifespan, the number and sex of dead, and number of censored (i.e., escapees, accidental deaths) events were recorded three times each week during transfer of flies to new food until all flies in each vial had died. To estimate fecundity, females were provided with fresh media each Monday and following a 24 hour egg laying period, these egg vials were collected on Tuesday and stored at −20 °C until processing. We continued this process each week until all females within a vial had died. We modified a protocol developed in Michael Rose’s laboratory by Larry G. Cabral at UC Irvine based on personal communication and video tutorials (Burke et al. 2016; Rose Lab Data 2012a,b) to filter eggs onto circular discs of black filter paper. Subsequently, discs bearing eggs were photographed using a Canon EOS Rebel TS*i* (Canon Inc, Japan) camera using a soft uniform lit custom studio placed in a dark room (see Ng’oma *et al*, 2018 for details). The following camera settings were used for imaging: exposure time 1/25 seconds, aperture F5.6, and ISO 100. Image files were uploaded to a computing system and immediately inspected with ImageJ (Schindelin et al, 2012, Rueden et al 2017) for quality. A detailed description of our method can be found in Ngoma et al (2018).

### Statistical analysis

All following statistical analyses were performed in R (ver. 3.5; R Core Team 2018), and all code is available online (https://github.com/EGKingLab/h2lifespan).

### Survival estimates

We converted daily counts of dead and censored individuals to individual events at a given age to perform survival analysis. Using the R package survival (ver. 2.38; (Therneau and Grambsch, 2000; Therneau, 2015), we used the Kaplan-Meier estimator (Kaplan and Meier, 1958) to estimate survival in each diet:

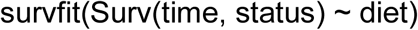

where time is time (days), status is alive/dead, diet is diet treatment (C, DR, HS) in order to visualize survivorship curves in each diet.

To assess statistical differences between diets accounting for family membership, we used hierarchical Bayesian inference to test for the effects of sire, dam nested in sire, vial nested in dam, and treatment using a set of six nested models, which tested variously for effects of diet treatment, sire, dam, and vial (Table 1). We used the brms interface (Bürkner, 2017) to the statistical modeling language stan (Carpenter *et al.*, 2017), which samples posterior estimates of modeled parameters using Hamiltonian Monte Carlo. Lifespan was modeled with a Weibull distribution, which is commonly used to analyze time to failure (in this case, fly death). We used broad priors for the Weibull shape parameter (Γ (0.01, 0.01)), Student’s *t*(3, 4, 10) for the intercept, and Student’s t(3, 0, 10) for the standard deviation, which were quickly overwhelmed by the large amount of data (7486 observations). Models were compared using leave-one-out (loo) cross-validation and loo model weights (Vehtari *et al.*, 2017; Yao *et al.*, 2018), the latter which assigns relative out of sample predictive ability to each model compared. Posterior estimates of DR and HS diet treatments were compared to the C diet treatment via credible intervals. We sampled each model for 2000 iterations with 50% discarded for warm-up and 4 replicate chains, to yield approximately 4000 total samples. We found that due to the large amount of data, this sampling approach was adequate. Adequate sampling was determined by 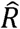 values of 1 (Gelman and Rubin, 1992; Brooks and Gelman, 1998).

**Table 1.** Comparison of models of lifespan, early fecundity, and total fecundity across diet treatments. Models 1 and 2 do not include any aspects of pedigree, and models 3-6 use a hierarchical model. Terms in parentheses are random effects. All random effects were “intercept” terms, allowing separate intercepts for each level of sire ID, dam ID nested in sire ID, or sire ID nested in diet treatment.

|  | **Model** | **Lifespan** | **Early**<br>**Fecundity** | **Total**<br>**Fecundity** |
| --- | --- | --- | --- | --- |
| 1 | ~ Intercept | 5% | 3.1% | 1.1% |
| 2 | ~ Diet | 0% | 10.3% | 0% |
| 3 | ~ Diet + (Sire ID) | 0% | 0.0% | 2.1% |
| 4 | ~ Diet + (Dam ID by Sire ID) + [Vial ID by Dam ID]<sup>†</sup> | 0% | 86.6% | 19.4% |
| 5 | ~ Diet + (Sire ID) + (Dam ID by Sire ID) + [Vial ID by Dam ID]<sup>†</sup> | 0% | 0% | 0% |
| 6 | ~ Diet + (Sire ID) + (Dam ID by Sire ID) + (Sire ID by Diet) + [Vial ID by Dam ID]<sup>†</sup> | 95.1% | 0% | 77.4% |

<sup>†</sup>Only the models for lifespan included this vial effect because the response variable for fecundity is a vial average.

### Fecundity estimates

We used a high through-put method to extract egg counts from images by constructing an optimal predictive model. To accomplish this, we took advantage of the simple relationship between number of eggs present and the amount of white area on a thresholded image of disc and used a set of hand-counted images to optimize the model. We were able to determine the optimal number of hand-counted images, select an appropriate threshold value, and assess model performance across a range of parameters. This method performs very well, with a 0.88 correlation between the model predicted egg counts and hand counted egg counts. A detailed account of our method is presented elsewhere (Ng’oma et al 2018).

We focused on two fecundity measurements from our weekly fecundity measures. We calculated the number of eggs per female for each of our weekly measures by dividing by the number of females alive in each vial. We then obtained an estimate of total fecundity per female by simply summing across weeks (hereafter referred to as total fecundity). We note that this is not strictly a measure of lifetime fecundity as we measured fecundity only once per week, though we would expect this estimate to be highly correlated with lifetime fecundity. Second, we considered a snapshot of early life fecundity by choosing the time point closest to 5 days post-eclosion (hereafter referred to as early fecundity). The actual age of the females varies slightly as the vials were set up over several days, but our fecundity measurements always took place on a 24 hour period beginning on Monday.

We tested for the effects of sire, dam nested in sire, and treatment using a set of six nested hierarchical (mixed) linear models (Gelman and Hill, 2007), which tested variously for effects of diet treatment, sire and dam, identical to the models used to test survival estimates but without the vial effect given our measurements for fecundity are averages per vial (Table 1). Sampling was done on zero-centered values for early life and total fecundity using Hamiltonian Monte Carlo using the statistical modeling language stan (Carpenter *et al.*, 2017), via the rstanarm interface (ver. 2.13.1; (Stan Development Team, 2016). We used mildly regularizing priors: Normal(0, 10) for intercepts, Normal(0, 10) for diet treatment parameters, and Cauchy(0, 1) for variances. Priors for covariance matrices were set to 1 for regularization, concentration, shape, and scale (i.e., decov(1, 1, 1, 1)). Models were sampled for 20,000 iterations, with 10,000 discarded for burnin. Adequate sampling was assessed by 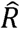, and models were compared using leave-one-out (loo) cross-validation and loo model weights as described above.

### Heritability in single diets

To estimate variance components, heritability, and phenotypic correlations, we used MCMCglmm (v. 2.24; Hadfield, 2010), a Bayesian inference package for R (v. 3.5; R Core Team, 2016) to fit variations of the ‘animal model’ (Kruuk, 2004; Wilson *et al.*, 2010; Ingleby *et al.*, 2013). Using the animal model we first inferred, in each diet separately, the additive genetic effect (*V*_*A*_) from pedigree information where the identities of sire and dam are included in the random effect

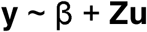

where **y** is the vector of phenotypes (lifespan, early fecundity, or total fecundity), β is the mean phenotype, **Z** includes information about sire and dam identity (the “animal” term), and **u** is the parameter vector that is estimated for the random effect, distributed as Normal(0, σ^2^). We did not include vial in order to estimate vial effects as with a half-sibling design in which half-siblings do not share the same vial, these effects will not influence the estimation of the additive genetic variance. We used weakly informative priors: *V* = 1, *v* = 0.002 for analysis of lifespan, although analyses were robust when more informative priors were tested. Similarly, we used weakly informative, parameter expanded, priors for analyses of fecundity. Parameter expanded priors, which improve prior distributions of variance components, can improve sampling in some cases (Hadfield, 2018). The models were sampled for 2 × 10^6^ iterations, discarding the first 1.5 × 10^4^ iterations as burnin, with a thinning interval of 50 iterations. These sampling parameters yielded effective sample size >10,000. We evaluated the models for convergence and autocorrelation of the samples, ensuring autocorrelation of <0.1 for the first lag, and absence of trend in the trace plots of the mean β (i.e., intercept term), additive variance *V*_*A*_, and residual *V*_*R*_ (the later two denoted in the output as ‘animal’ and ‘units’, respectively).

### Genetic correlations between diets

A single phenotype that is measured in multiple environments (here diets) may be regarded as different phenotypes in those environments (Falconer, 1952; Lynch and Walsh, 1998). Based on this premise, we computed genetic correlations for phenotype-diet combinations by fitting a Bayesian multivariate animal model:

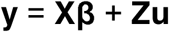

where **y** is a matrix of phenotypes (C, DR, HS), β is a vector of trait (**X**) means (with no intercept term), **Z** includes information about sire and dam identity (the “animal” term), and **u** is the parameter vector that is estimated for the random effect. Among the random effects, covariance was free to vary (i.e., ‘random = ∼us(trait):animal’) with one restricting the variance to zero (i.e., ‘random = ∼idh(trait):animal’). If family genetic effects remain the same or change in proportion between diets, then the genetic correlation of family members between diets is 1. Genetic correlations that are credibly smaller than 1 indicate a significant genotype × diet interaction. We therefore compared the model above with a model in which the correlation between diets is constrained to be 1 to test if our estimated genetic correlations were significantly different from 1. In bivariate models, we tested several slightly stronger variations of priors (*V* = diag(3), *v* = 1.002). As above, we used a parameter expanded prior for fecundity. Model were run for 6.5 × 10^6^ iterations, with 5 × 10^4^ discarded as burning, with thinning every 500 iterations. Twelve chains were run in parallel, yielding >10,000 samples total.

## Results

### Phenotypic response to dietary treatment Lifespan

We used a split family design, splitting offspring from families into three different diets. Relative to the control diet, median survival was 24% lower on high sugar (HS: 48 days vs. C: 63 days) and 8% higher on DR diet (68 days) (Fig. 2b). Lifespan trajectories began to diverge early, from about 25 days post oviposition and remained diverged until under 10% survivorship (Fig. 2b, Table S3). Individual sire family responses to diet are presented in Fig. S1.

**Figure 2:**
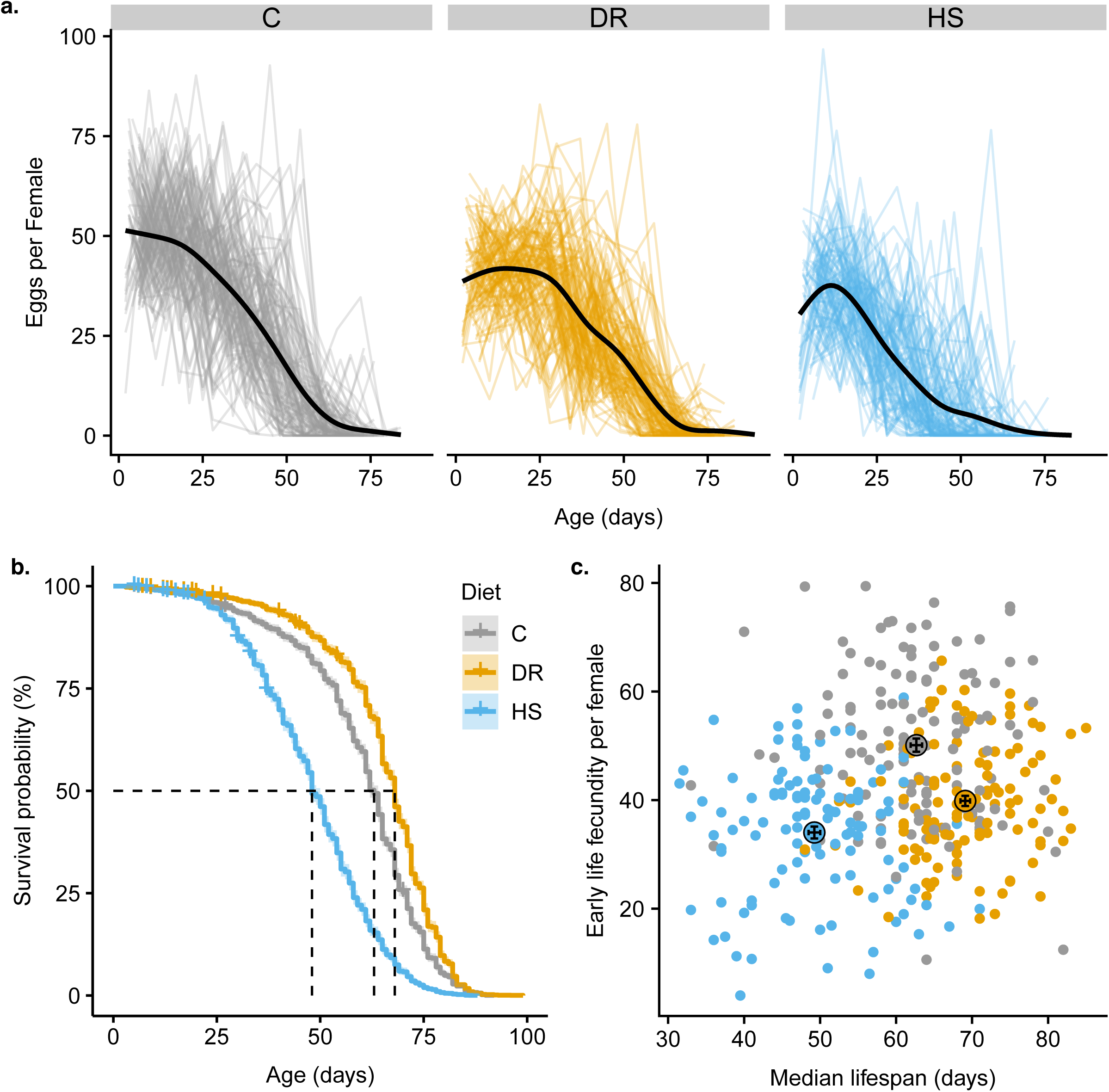
Phenotypic response to diet treatment. The different colors represent different diet treatments (grey = C, orange = DR, blue = HS). **a.** Fecundity per female sampled over a 24 hour period once per week over lifespan. A smooth (loess) fit is shown for each treatment. Individual lines represent each replicate vial. **b.** Survival probability versus age in each of the three diet treatments. Median survival (50% survival probability) is shown with the dotted line for each diet. Censored events are marked with a + sign on each survivorship curve. **c.** The relationship between early life fecundity (24 hour sample period closest to 5 days post-eclosion) and median lifespan for each vial. The average values for both within each treatment are shown by the large point outlined in black. Error bars are the mean +/-1 standard error. Note that the error bars all fit within the size of the point.

We compared six nested models of survival, incorporating diet treatment and various aspects of relatedness and compared these models using leave-one-out model weighting (Table 1). The model including sire, dam nested in sire, vial nested in dam, diet, and a sire by diet interaction has the overwhelming model support (>95% of model weight), indicating a significant effect of diet, the presence of significant genetic variance for survival within the diet treatments, and significant genetic variation for the response to diet. Specifically, the results of this favored model show that, compared to the C diet group, individuals in the DR group were credibly less likely to die (99% credible interval of the difference between DR and C = 0.4 - 0.12) and individuals in the HS treatment group were credibly more likely to die (99% credible interval of the difference between HS and C = −0.21 - −0.13).

### Fecundity

Fecundity was estimated once per week in the same treatment setups from which lifespan was recorded in order to compare the two phenotypes in the same set of flies. As expected, in all treatments, fecundity declined with age (Fig. 2a). We used a Bayesian model comparison approach to analyze the effects of diet and family on total fecundity and early fecundity (Table 1). This analysis revealed a strong effect of diet for both total fecundity and early fecundity, as models including diet were strongly preferred over a model without diet. For both measures of fecundity, the highest values were in the C diet and the lowest in the HS diet (Fig. 2a,c). There was a credible effect of family for both fecundity measures, indicating significant genetic variation for fecundity. For early fecundity, the model including only dam nested within sire was favored heavily. For total fecundity, the model that included sire, dam nested in sire, and the interaction between sire and diet was favored over the other models, indicating genetic variation for the response to diet in addition to genetic variation for fecundity.

### Estimates of quantitative genetic parameters

We have a high sample size for lifespan with data for up to 48 individuals spread across two vials for each dam family in each diet. While these same individuals contributed to our measurements of weekly fecundity, we have a single measurement for each vial of females at any given time point and do not have measurements per each individual, as measuring fecundity for each individual female was not feasible for this study. Thus, for estimating quantitative genetic parameters, our replication for fecundity is low (2 per family per diet). The results of the above models reflect the expected uncertainty from this low replication with wide posterior distributions. Thus, while the models we fit above demonstrate the presence of significant genetic variation, our estimates of heritability values have high uncertainty and wide credible intervals. Therefore, we focus on discussing our results for lifespan below and results for fecundity can be found in Fig. S2.

### Heritability of lifespan within diets

We estimated the heritabilities of our phenotypes separately in each diet treatment using a Bayesian approach (see Methods). Our heritability estimates for lifespan are moderately high, ranging from 0.31 to 0.47 (Figure 3a). Heritability (*h*^*2*^) and 95% HPDI (highest posterior density interval) for each treatment were: C *h*^*2*^ 0.47 (0.34 – 0.61), HS *h*^*2*^ 0.37 (0.25 – 0.50), DR: *h*^*2*^ 0.31 (0.21 – 0.43). 95% HPDIs of pairwise differences of posterior estimates of heritability did not differ credibly from 0: C-DR = - 0.02 – 0.34, C-HS = −0.09 – 0.29, and HS-DR = −0.11 – 0.23.

**Figure 3:**
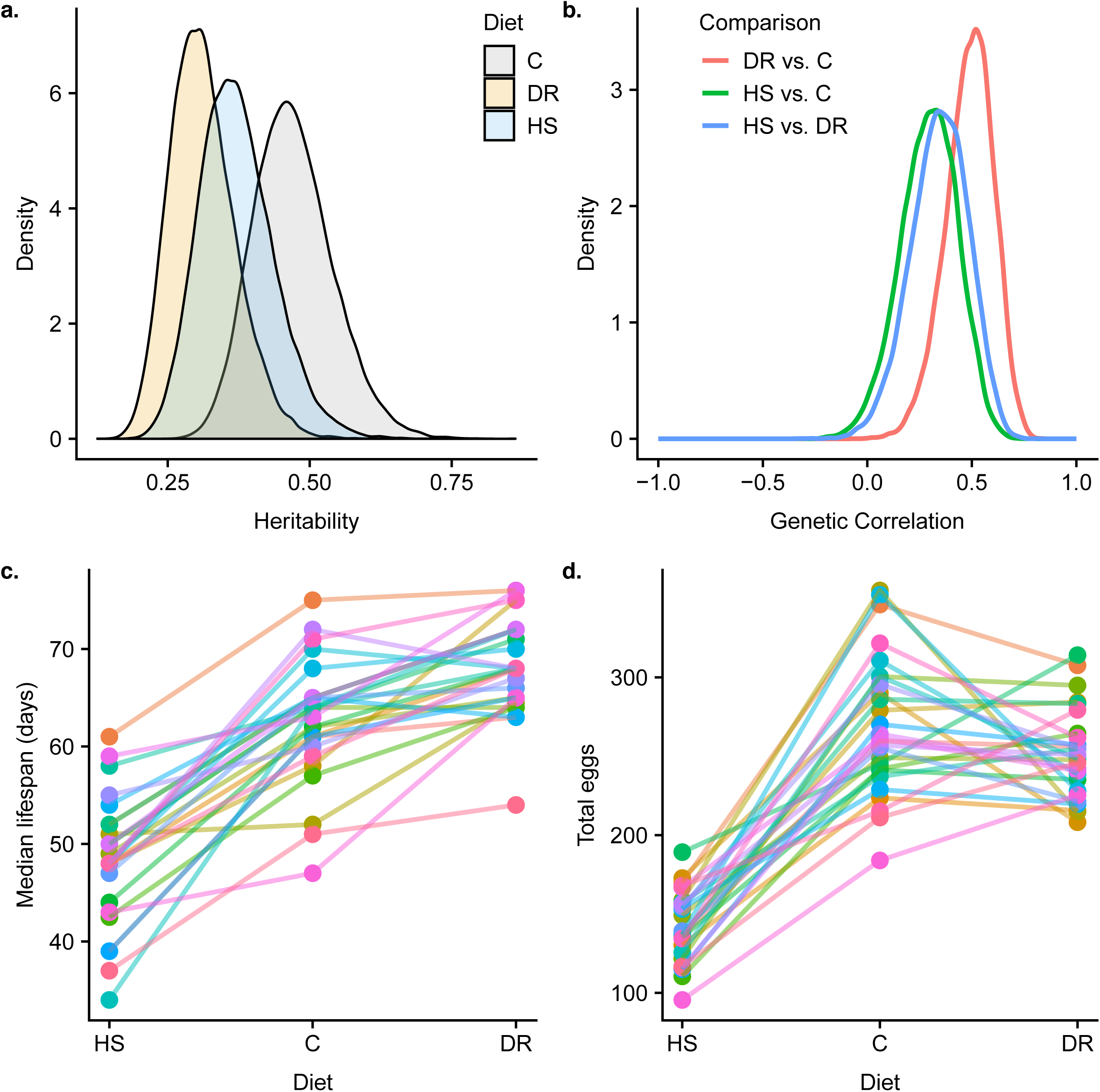
**a.** Density plots of the estimated posterior probabilities for lifespan heritability within each diet. **b.** Density plots of the estimated posterior probabilities for the genetic correlation between lifespan in pairs of diets. **c.** Sire by environment interaction plot for median lifespan showing the median lifespan for each sire family in each of the three diets. Each line represents one sire family. **d.** Sire by environment interaction plot for total egg count per female (summed across weekly measures) showing the median lifespan for each sire family in each of the three diets. Each line represents one sire family.

### Correlation of lifespan across dietary treatments

To determine if there is substantial genetic variation for the response to the environment (GEI), we fit a multivariate form of the animal model, which considers lifespan within each diet as a separate phenotype (Falconer, 1952; Lynch and Walsh, 1998; Wilson *et al.*, 2010). When the genetic correlation between environments is less than one, it indicates significant genetically based differences in the response to the environment, or a significant GEI. In all pairwise diet comparisons, our estimates of the genetic correlations are credibly less than one, indicating substantial genetically based variation in the response to diet (Fig. 3b). Genetic correlations (*r*_*g*_) and HPDIs were: DR vs. C: *r*_*g*_ 0.48 (0.34 – 0.61), HS vs. C: *r*_*g*_ 0.29 (0.21 – 0.43), HS vs. DR: *r*_*g*_ 0.34 (0.25 – 0.50. This variability is also apparent when examining the variation among sire families in their response to diet for both lifespan and total fecundity (Fig. 3c & d). In addition, the model comparisons we employed for survival and total fecundity both favored models including a sire by diet interaction, which is consistent with the presence of a significant GEI. Subsequent analysis of differences of posterior estimates of genetic correlations did not reveal credible differences: HS vs. C - DR vs. C = −0.51 – 0.12, HS vs. C - HS vs. DR = −0.35 – 0.24, and DR vs. C - HS vs. DR = −0.19 - 0.47, indicating similarities in genetic correlations among treatment pairs.

## Discussion

In this study, we used an admixed multiparent population of *D. melanogaster* to characterize the quantitative genetics of fitness components in different nutritional conditions. Our study is one of only a few studies that estimate both lifespan and lifetime fecundity across multiple diets in the same families, providing a comprehensive picture of the interplay between the genetic basis of important life history traits and the nutritional environment. We are able to show, not only that these traits harbor substantial genetic variation both within diets and in the response to diet, but we also provide concrete estimates of both the narrow-sense heritability of lifespan in multiple diets and the genetic correlation of lifespan between diets. These results have important implications for our understanding of the evolution of these traits in wild populations and for strategies to uncover the genetic basis of these traits, which we discuss below.

### Evolutionary potential of the coordination between nutrition and fitness components

We characterized the effects of three different diets, a dietary restriction diet (DR), a control diet (C), and a high sugar diet (HS). Dietary restriction has been studied extensively in *D. melanogaster* and other metazoans (e.g., Chippindale *et al.*, 1993a; Good and Tatar, 2001; Vrtílek and Reichard, 2014; reviewed in: Sohal and Weindruch, 1996; Kenyon, 2005; Kirkwood and Shanley, 2005; Piper *et al.*, 2011; Tatar, 2011; Tatar *et al.*, 2014). A general finding of these studies is that reduced nutrient intake without starvation results in lifespan extension, and that this effect is often coupled with reduced reproductive output, though there are exceptions to this pattern (e.g., Kaitala, 1991; Weindruch *et al.*, 1995; Kirk *et al.*, 2001; Stelzer, 2001; Cooper *et al.*, 2004; reviewed in Ng’oma *et al*., 2017). Even within species that tend to show this effect such as *D. melanogaster*, there is variability in this pattern among different genotypes. For example, Stanley *et al.* (2017) found variation among inbred lines for the amount of lifespan extension, with some lines showing little to no extension in a DR diet, and a few showing reduced lifespan in a DR diet, a result that has also been shown in mice (Rikke *et al.*, 2003). Unlike for DR, the effects of a high sugar diet have not been studied as extensively, though recently some studies have shown flies reared on high sugar diets as larvae show obesity-like phenotypes (Musselman *et al.*, 2011; Rovenko *et al.*, 2015), and adults reared on high sugar diets have elevated triglyceride levels, reduced lifespan, and reduced fecundity (Skorupa *et al.*, 2008), a pattern that is consistent with our findings. In these studies, “high sugar” refers to a high concentration of sugar relative to the volume of the diet, and that is how we also use the term to describe our high sugar diet. However, nutritional geometry approaches focus on the protein to carbohydrate ratio of the diet, and thus “high sugar” diets in this case would correspond to a diet with a low P:C ratio. In this context, our HS diet, with a yeast-sugar ratio of about 1:2 resulted in reduction of both lifespan and fecundity. Past studies (Skorupa *et al*, 2008, Edgecomb *et al*, 1994) observed that flies on a diet richer in carbohydrate tended to feed more, and that imbalance in nutrient intake resulted in shorter lifespan. Other studies including Lee *et al* (2008) and Jensen *et al*, (2015), on the other hand, found lifespan and lifetime fecundity maximized at much lower P:C ratios (1:16 and 1:4, respectively). Apparent incosistence of our results (and studies cited above) with these studies that measure specific nutrient content likely reflect diet component composition relative to volume. We note, however, that our diets use agar to form a semi-solid media while these studies use a liquid diet, and that our flies are maintained under 24 hour light conditions, both of which may also lead to differences in the patterns observed.

These types of phenotypic patterns in different diet conditions have been interpreted in different ways. The widespread observation of lifespan extension observed in response to dietary restriction in many taxa has led to competing hypotheses for this pattern. It is often argued that this pattern is the result of natural selection acting in environments with fluctuating resources where it should be adaptive to increase allocation of nutrient resources to the soma when nutrients are limiting, however it is also possible it is the result of an unavoidable physiological constraint (Holliday, 1989; Kirkwood and Shanley 2000; Kirkwood and Shanley, 2005).

Additionally, studies employing high sugar and high fat diets often view the results of those studies in the context of human populations where it is hypothesized the negative health consequences of a Westernized diet result from a mismatch between that diet and the diet humans adapted to in the past over the majority of their evolutionary history (Neel, 1962; Wells, 2009), though this hypothesis has also been challenged (Speakman, 2008). Lately, a number of studies have challenged the resource allocation framework to explain life history trade-offs (see Barnes *et al*, 2006; Flatt *et al*, 2008; Adler *et al*, 2013). These studies (and others) suggest that the decoupling of lifespan from fecundity that is sometimes observed, relates to hormonal signaling rather than literal resource allocation. However, the works of Zhao and Zera (2006), Zera and Zhao (2006), and Zera (2005) demonstrate the biochemical basis of life history trade-offs (reviewed in Ng’oma et al 2017). In general, assessing the potential for phenotypic effects of different diets to be adaptive patterns or to be highly constrained physiological responses requires an understanding of the underlying genetic variation for the response to diet in order to understand if this response has the potential to evolve.

Here, we showed there is heritable variation for both phenotypes within diets, and for the response of phenotypes to diet (Table 1; Fig. 3a,b). Previous mapping studies in *D. melnanogaster* have identified several QTL influencing life history traits (reviewed in Paaby and Schmidt, 2009), including mapping QTL influencing changes in gene expression and lifespan in different diets (Stanley et al. 2017). However, like other complex traits, these QTL fail to explain most of the underlying genetic variance. Our results support the hypothesis that these phenotypes are influenced by many genes and quantify the amount of variation among individuals attributable to genetic variation in this population. While many studies have demonstrated GEI for different traits (Reed *et al.*, 2010; Qi *et al.*, 2012; Ingleby *et al.*, 2013), few studies have estimated GEI specifically for life history traits in response to diet, and not all previous studies of diet have found significant GEI. For example, King *et al.* (2011) estimated quantitative genetic parameters for allocation to flight capability and to reproduction in the sand cricket, *Gryllus firmus.* These authors found no GEI for these traits, with all genotypes allocating more to reproduction when food levels increase, suggesting important evolutionary constraints can exist for the response to diet and that significant GEI is not a foregone conclusion. Our results show that this constraint does not exist for this population of *D. melanogaster* and that the response to diet may readily evolve in response to selection. The founder lines of the DSPR have a global distribution and were derived from wild-caught flies and thus, our population encompases some of the genetic variation that exists among natural populations of *D. melanogaster.* It is plausible, therefore, that the diet dependent effects observed in this study and others represent adaptive patterns that have evolved. Some recent experimental evolution studies exposing flies to constant DR, standard, and high protein diets have shown mixed and sex-specific results in which females exposed to high protein diet evolved increased fecundity but decreased survival on standard and DR diets (Zajitschek et al 2016), while males on DR had greater fitness compared to those on standard and high diets (Zajitschek et al 2018). These studies establish that the responses of fecundity and lifespan to diet do evolve, but the emergent patterns do not support the typical hypothesis that selection on differential resource allocation is driving the pattern. It remains a challenge to interpret how the patterns we observe in these kinds of laboratory studies represent the nutritional conditions experienced by flies in the wild and by what mechanisms those conditions have driven the evolution of different life history strategies. It would be interesting to see if future studies will also find evoltuionary potential for the response to changes in the P:C ratio.

### Prospects for multiparent population approaches

One of the most fundamental goals in biology is to understand the genetic basis of complex phenotypes. This goal has proved to be quite challenging. For most traits, some of which have been the focus of study for many years, the causative genetic variants that have been discovered explain only a small percentage of the trait heritability (for reviews see McCarthy *et al.*, 2008; Manolio *et al.*, 2009; Rockman, 2012; Visscher *et al.*, 2012). For traits that are expected to be highly polygenic, such as lifespan and fecundity, estimating quantitative genetic parameters as we have done in this study, can be as informative about the genetic basis of a trait and its evolutionary potential as studies aimed at uncovering specific genetic variants.

In recent years, multiparent populations, in which multiple inbred founder lines are crossed for a number of generations to create a recombinant mapping population, have increasingly been employed in studies aiming to identify the causative genetic variants underlying complex traits (e.g. (Kover *et al.*, 2009; McMullen *et al.*, 2009; Huang *et al.*, 2011; Aylor *et al.*, 2011; Threadgill and Churchill, 2012; King *et al.*, 2012; King, *et al.*, 2012; Cubillos *et al.*, 2013); reviewed in (Long *et al.*, 2014; de Koning and McIntyre, 2017). One reason for employing a mapping population consisting of multiple parent lines is the inclusion of additional genetic variation when compared to traditional two-line QTL mapping panels. Even with multiple parent lines contributing, a potential criticism of these kinds of approaches when compared to mapping approaches that sample from natural populations is that they contain limited amounts of genetic variation. The DSPR population used in this study was started from 8 inbred parent lines. Therefore, while every individual in our admixed population has a unique mosaic genome, at any given position, there are only 8 possible haplotypes. Despite this simplicity, we find moderate heritabilities for lifespan in all diets (ranging from ∼0.3 - 0.5) that are comparable to estimates from natural populations. A meta-analysis of heritability estimates from natural populations found life history traits typically have values around 0.25 and values higher than 0.3 are not infrequent (Rose and Charlesworth, 1981; Mousseau and Roff, 1987; Rikke *et al.*, 2010). We also found strong evidence for a genotype by diet interaction for lifespan, indicating there is also segregating genetic variation for the response to diet. While our evidence of a genotype by diet interaction is not as strong for fecundity, our model comparison for total fecundity supports a model including this interaction.

In addition to their utility for QTL mapping, multiparent populations have also been used in evolve and resequence approaches (e.g. Turner and Miller, 2012; Burke *et al.*, 2014a,b) to identify the underlying genetic basis of complex traits. These studies impose selection and track the resulting changes in phenotype and genotype. This approach is potentially very powerful because 1) entire haplotypes can be tracked over evolutionary time, 2) the simplified genetic architecture with only a few potential haplotypes at any given location may lead to a more repeatable response to selection increasing the likelihood of identifying causative loci, and 3) it may allow for validation of QTL identified in mapping studies using these same populations. A potential downside to such an approach is if the relatively simple genetic architecture of such a population results in limited genetic variation for the trait of interest, thereby constraining evolution. Our study shows there is substantial genetic variation for our focal phenotypes in our admixed DSPR population, which establishes this approach as a viable strategy for investigating both how a focal trait evolves and the underlying genetic basis. It is one of only a few studies to quantify quantitative genetic parameters for an outbred multiparent population, providing evidence that segregating genetic variation is not limiting in these populations. In general, MPPs have great potential to advance our understanding of the genetic basis of complex traits via these kinds of multi-pronged approaches.

## Acknowledgments

We received helpful suggestions from Larry Cabral on how to establish our fecundity protocol. We had insightful discussions with Ian Dworkin on fitting the animal model in the Bayesian framework. Stuart Macdonald provided the DSPR RILs used to start our experimental colony. Elizabeth Lopresti, Michael Reed, Osvaldo Enriquez, Anna Perinchery and Kyla Winford helped with fly husbandry, experimental set-ups, collection and entry of data. This work was supported by NIH grant R01 GM117135 to Elizabeth G. King, The University of Missouri, and a University of Missouri Research Board Grant.

## Data availability

Raw data, including lifespan, fecundity, and image files, may be retrieved from Zenodo at http://doi.org/10.5281/zenodo.1285237. Scripts to reproduce all analyses are available at GitHub: https://github.com/EGKingLab/h2lifespan. Supplementary information is available at www.nature.com/hdy/

